# Hydropeaking intensity alters riparian seed bank composition and self-restoration potential

**DOI:** 10.64898/2026.08.15.745034

**Authors:** Emil Nordström, Sergey Rosbakh, Jacqueline H. T. Hoppenreijs

## Abstract

Flow regulation for hydropower production affects stream ecosystems through decreased connectivity and changed timing and magnitude of flows. Hydropeaking, a form of regulation in which infrequent large peak flows are replaced with frequent small peaks, increases riparian erosion and causes water and drought stress for riparian vegetation. Hydropeaking is likely to affect soil seed bank (SSB) formation and composition, while SSBs are important sources of self-restoration should a system’s flow regulation be relaxed. We tested how hydropeaking intensity affects the size and composition of SSBs, including the functionally important group of large graminoids, and by calculating Ellenberg values for Moisture, Light and Soil disturbance. SSB samples were taken at 15 riparian zones across central and northern Sweden. Each site was regulated, but sites differed in their hydropeaking intensities. SSBs were subjected to a seedling emergence experiment, from which over 700 seedlings from 53 taxa emerged. We found that hydropeaking intensity affects the composition of soil seed banks on multiple levels. Seedling density was negatively correlated with hydropeaking intensity at the sites where samples were taken. SSB richness varied (two to eighteen species per site) and was not affected by hydropeaking intensity. The proportion of large graminoids in the seed bank showed a near-significant decrease with increasing hydropeaking intensity, and community-weighted means for Moisture, Light and Soil disturbance increased (non-significantly) with increasing intensity. Our results suggest that riparian SSBs, should flow regulation be relaxed or ceased, are not sufficient for self-restoration of functional riparian vegetation. Seeds of large graminoids and species that are tolerant to drought in the germination stage are less present in riparian SSBs of heavily-regulated streams. Supply of seeds of these groups, or even planting, may need to be considered when changes in flow management are implemented.

**Highlights:**

- Hydropeaking negatively affects riparian soil seed banks (SSBs) in Sweden
- SSB size slightly decreases with hydropeaking intensity, but richness does not change
- The proportion of large graminoid seeds in SSBs decreases with hydropeaking intensity
- Riparian SSBs from less-impacted sites have most potential for self-restoration

## Introduction

Rivers and riparian zones are crucial for human and natural sustenance, but globally fragmented and degraded by human activities (Dudgeon, 2019). Located on the interface of terrestrial and freshwater ecosystems, riparian zones maintain water quality through nutrient cycling, physical buffering and temperature regulation (Arthington et al., 2010). Freshwater and riparian biodiversity are closely interconnected, as many species depend on both habitats for different phases of their life cycles, and because their food webs are intertwined (Nakano & Murakami, 2001). Riparian communities are shaped by the river’s flow regime and its interaction with local and regional geomorphology, which results in a dynamic mosaic of vegetation (Corenblit et al., 2007; Harris et al., 2026). Laterally, riparian vegetation shows a zonation pattern with smaller, more flexible and resilient species close to the water line and larger, sturdier and more resistant species further away from the water, towards upland areas (Figure 1). Under the pressure of human activities, such as reduction of peak flow, and through the changing climate, riparian zones become narrower or disappear completely, and riparian vegetation can become less species-rich or develop a different functional composition (Hoppenreijs et al., 2022; Jansson et al., 2019).

**Figure 1.**
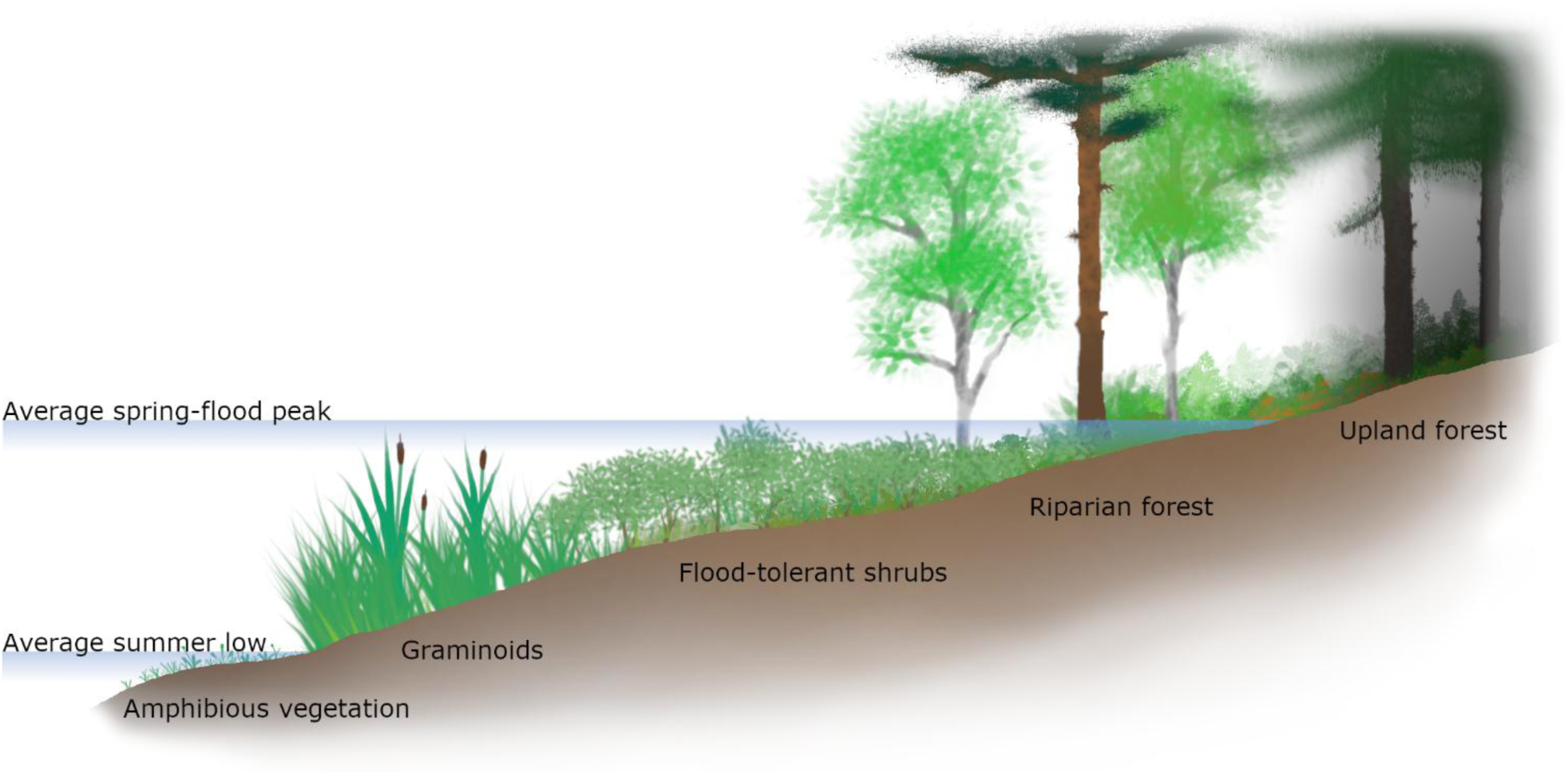
Generalized view of a riparian zone, showing the zonation in vegetation and their relation to the water level. Modified from Ström et al. (2012).

Hydropower is one of these human activities and, while being an increasingly important source of renewable energy, drives freshwater and riparian degradation locally and regionally (He et al., 2024). Water releases from hydropower dams rarely match the timing, magnitude or frequency of natural peak flows, and the dams themselves form barriers for in-stream matter and organisms. Hydropower thus changes connectivity along one or multiple dimensions of the ecosystem (Baird et al., 2024; Ward, 1989). More intensive forms of hydropower production such as hydropeaking, in which smaller amounts of water are released frequently, not only change the timing of high- and low-flow events but also intensify cycles of flooding and drying. This results in mechanical and water stress for aquatic and riparian vegetation, causing changed germination, establishment and growth among riparian plants (Bejarano et al., 2018). Differences in functional composition of regulated (hydropeaking and other forms) rivers is widely reported (e.g., Jansson et al., 2000; Lytle et al., 2017), although the direction and extent of change may vary between systems (Lozanovska et al., 2020). The combination of functional shifts and altered reproductive ability of species (Bejarano et al., 2018), will result in different SSB composition. Riparian SSBs can undergo further change as the intense pulses of hydropeaking also disrupt the natural sedimentation and erosion regime (Bejarano et al., 2018). Hydropeaking leads to increased erosion (Nordström et al., submitted), which can remove surface soil layers and causes physical loss of SSBs. Simultaneously, the changes in high-flow timing could trigger germination for some species at unfavourable times of the year (Bejarano et al., 2018), causing SSB impoverishment. Thus, hydropeaking can change riparian vegetation and SSBs in various ways.

Hydropower production thus negatively affects riparian zones in multiple ways, but researchers, industry and governments explore methods for more sustainable exploitation of river flows (Nyqvist et al., 2025; Widén et al., 2022). Examples of such methods are minimal flows, in which river sections that fall dry under traditional flow regulation are guaranteed a continuous minimum amount of water flow, and environmental flows, in which natural flow regimes are mimicked. Reducing pressure on ecosystems, however, does not guarantee ecological or functional recovery. For the system to be able to self-restore, sufficient propagules must be available, through dispersal or from the soil seed bank (SSB) (Florentine et al., 2023). In riparian zones, the SSB can be an important source of such propagules, especially if it contains viable seeds from conditions prior to regulation (Hall and Zetler, 2010; O’Donnell et al., 2015). Such propagules may germinate when environmental conditions become more favourable, such as under more sustainable hydropower production, but their abundance may have decreased under long-term pressure (Sarneel et al., 2024).

Here, we study the effect of flow regulation on the composition of riparian SSBs to obtain a better understanding of the potential for self-restoration of riparian zones. We compare SSBs from different regulated streams along a gradient in hydropeaking intensity. We expect hydropeaking to negatively affect riparian seedbanks, with (1) a negative correlation between hydropeaking intensity and seedling density, (2) a negative correlation between hydropeaking intensity and SSB richness, and (3) a negative correlation between hydropeaking intensity and the proportion of species belonging to the graminoid belt (Figure 1) in the SSB. Because of changed environmental filtering in the form of more intense flooding and drying cycles, we also expect (4) a shift towards species with preference for more light and soil disturbance, and less soil moisture.

## Methods

### Study system

We sampled SSBs at 15 sites along hydropower-regulated streams, spanning a latitudinal gradient of approximately 550 km across boreal and boreo-nemoral Sweden (Table 1). We included sites in both inland (up to an altitude of about 350 m) and coastal areas (Figure 2). All streams included in this study are snow-fed and originate in the Scandes and their sizes varied considerably, with yearly average flows ranging from below 10 m^3^/s to above 100 m^3^/s.

**Figure 2.**
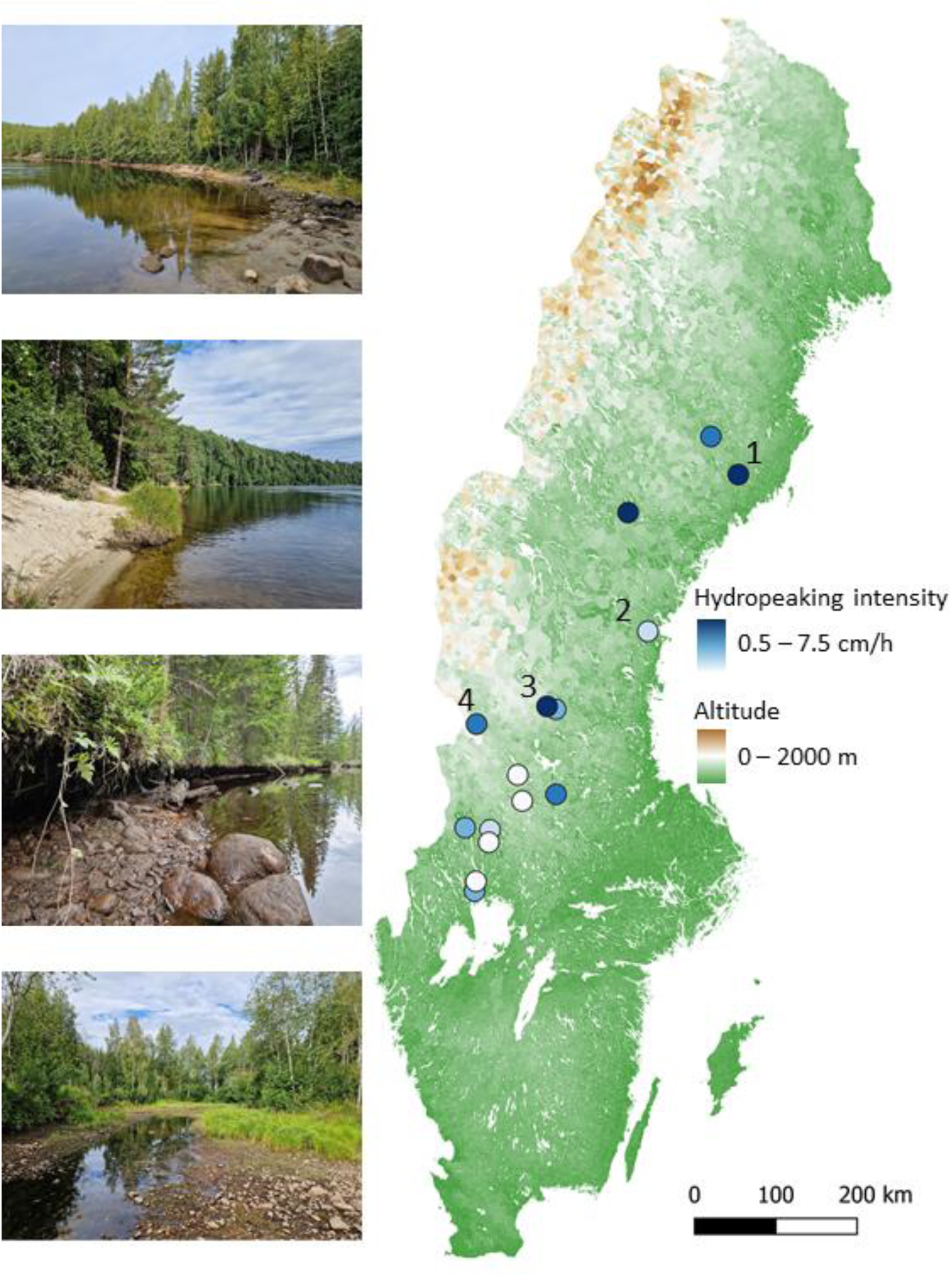
Examples of riparian zones and site map, note that the points are slightly displaced for visibility (see table 1 for precise site coordinates). Photos are (from the top): Harrsele (1 on the map), silty-sandy soil with mainly small hydrophytes as plant cover, intensive hydropeaking. Viforsen (2), a sandy site with very little hydropeaking. Vässinkoski (3), mixed substrate and considerable slope overhang, experiences occasional large fluctuations in water level from both hydropeaking and snowmelt. Horrmund (4), a high-altitude site with mixed substrate and extensive plant cover, hydrological regime is similar to Vässinkoski.

**Table 1.** Altitude is in meters above sea level, Distance corresponds to the distance (in m) from the sampling site to the dam or tunnel outlet, and hydropeaking intensity is measured as average hourly change in water level (cm/h).

| Site name | Coordinates<br>(WGS84) | Altitude (m above<br>sea level) | Distance from dam or<br>tunnel outlet (m) | Hydropeaking<br>intensity (cm/h) |
| --- | --- | --- | --- | --- |
| *Tuggen* | 64,4313, 18,9712 | 178 | 6750 | 3,8 |
| *Harrsele* | 63,9895, 19,6022 | 91 | 490 | 7,2 |
| *Långbjörn* | 63,6303, 16,7904 | 148 | 1030 | 5,7 |
| *Viforsen* | 62,3071, 17,1836 | 11 | 600 | 0,9 |
| *Vässinkoski* | 61,4924, 14,7854 | 344 | 1420 | 6,8 |
| *Noppikoski* | 61,4614, 14,9864 | 255 | 1400 | 2,3 |
| *Hormund* | 61,2830, 13,1613 | 351 | 290 | 4,5 |
| *Gävunda* | 60,7297, 14,1261 | 246 | 420 | 0,6 |
| *Lindbyn* | 60,5166, 14,9940 | 156 | 370 | 2,6 |
| *Eldforsen* | 60,4410, 14,2351 | 237 | 450 | 0,6 |
| *Röjdan* | 60,1314, 12,9709 | 82 | 1450 | 1,1 |
| *Edsforsen* | 60,0681, 13,5231 | 128 | 330 | 0,8 |
| *Skoga* | 60,0409, 13,5268 | 118 | 380 | 0,5 |
| *Frykfors* | 59,5245, 13,2522 | 55 | 540 | 0,8 |
| *Edsvalla* | 59,4429, 13,2197 | 44 | 450 | 1,1 |

For each site, we calculated hydropeaking intensity based on fluctuations in water level. Water depth was monitored continuously for one year prior to sampling with HOBO pressure loggers (model U20; Onset Computer Corporation, Bourne, MA) placed at all sites. Logger output was corrected for air pressure with loggers mounted above the water level and within 15 km of the site. As a measure of hydropeaking intensity, we used the average change in water level from one hour to the next (absolute values) which is similar to Richards-Baker index for flow data (Baker et al., 2004). We quantified the local magnitude of hydropeaking rather than frequently-used measures like distance from dam or hydropeaking volume measured at the dam, because this better accounts for local geomorphic variation (e.g., width of the river) and can be more directly related to mechanical and ecological effects at the affected sites (see inset photos in Figure 2). To our knowledge this method is, although similar to the Richards-Baker index, rarely used.

### Field sampling

Soil samples were taken in August-September of 2024. Samples were taken at three different elevations of the riparian zone (Figure 3), along two transects (apart from at Lindbyn, which had only one transect), in an area sheltered from the main flow. This approach ensures that local hydrogeomorphic variation is accounted for, which is necessary given that all sites are affected differently by their respective flow regimes. We estimated the elevations based on visible differences in vegetation (Figure 1), litter and substrate between the respective elevations in the riparian zone (which are linked to hydrological disturbance) and later verified these using the water level data. We collected 500 ml of soil from the first 5-7 cm of substrate, after removing any litter, in each sampling plot.

**Figure 3.**
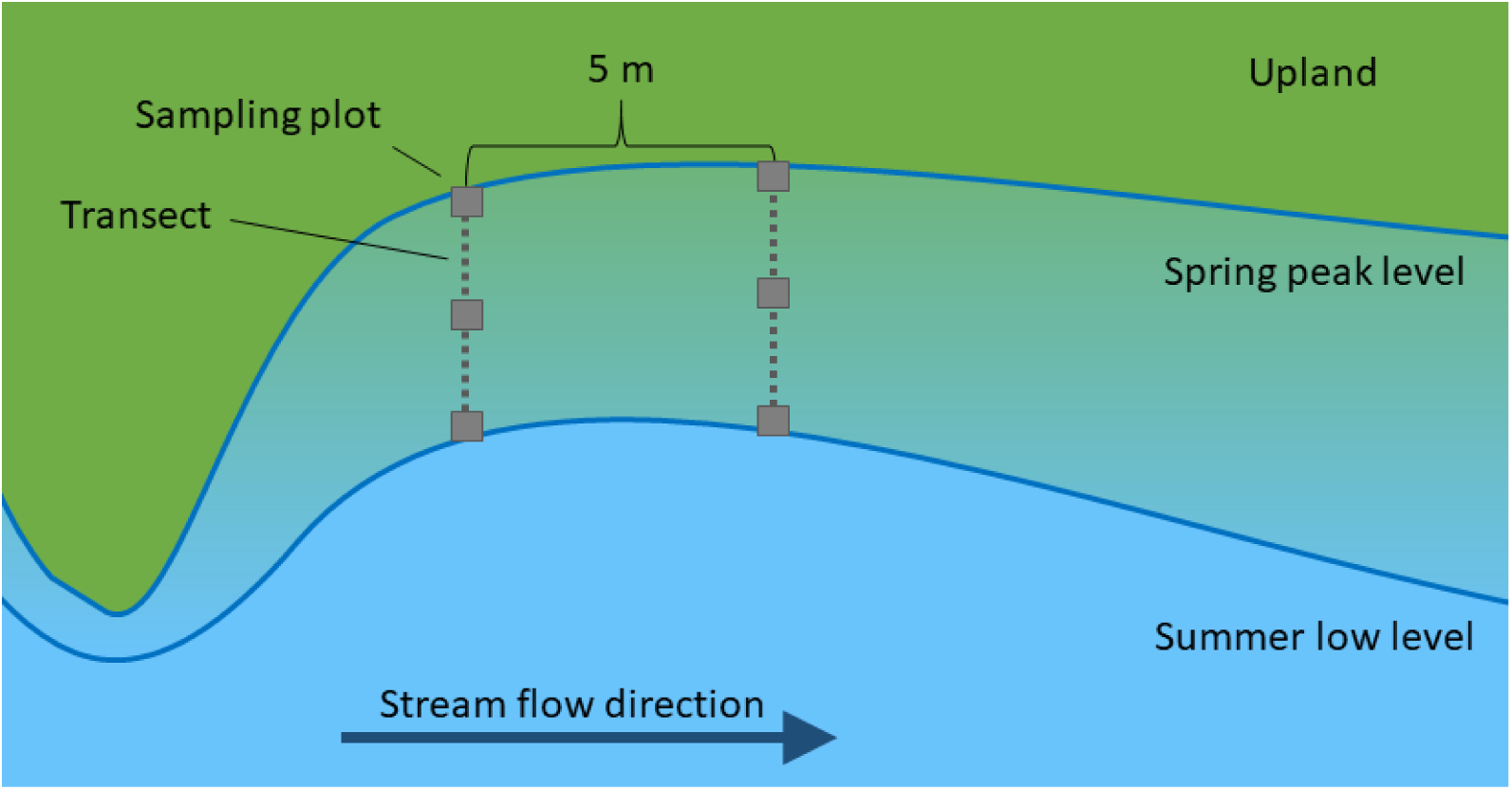
Placement of the sampling plots relative to the stream, with the yearly extremes in water levels indicated.

### Sample cultivation

The seed bank composition was identified through the seedling emergence method, applied to dried and sieved (at 4mm) samples (Ter Heerdt et al., 1996). The collected sediments were placed in plastic trays filled with a mix of equal parts pool filter sand (grain size 0.4-0.8 mm) and pond soil (30% clay, 40% “Sphagnum” (peat), and 30% sand), with 2 l of the mixture used per tray, filling it to a depth of approximately 2 cm. The trays were kept in a growing room under an average temperature and humidity of 18.4 °C and 74 % RH respectively with a photoperiod of 18/6 hours. The trays were divided into three sections, each with a replicate of 30 ml dry sample from a site, spread evenly across the section. Replicates were distributed randomly within and across trays.

We exposed the trays to three different treatments, designed to resemble different hydrological conditions normally occurring in the field, to allow species adapted to the full hydrological spectrum to germinate. The treatments were: mesic, daily waterlogging, and constant waterlogging (Figure 4). The mesic treatment was made to resemble near-upland conditions, and the trays sat on capillary mats and remained evenly moist but with no waterlogging. The daily waterlogging treatment was chosen to mimic conditions near the middle-elevation of the riparian zone under hydropeaking conditions, with the water level raised to the soil surface level each morning, and then allowed to drain over the course of the day (water level dropped to 0 after about 2 hours). The constantly waterlogged treatment corresponds to the conditions close to the stream, with the water level maintained near the soil surface throughout the study. Two replicates were included per site and treatment, apart from at two sites (Gävunda and Röjdan because of limited sample material). We used 30 trays in total, including two control sections per treatment (interspersed among the samples), to exclude the species whose seeds might be present in the substrate used for SSB cultivation.

**Figure 4.**
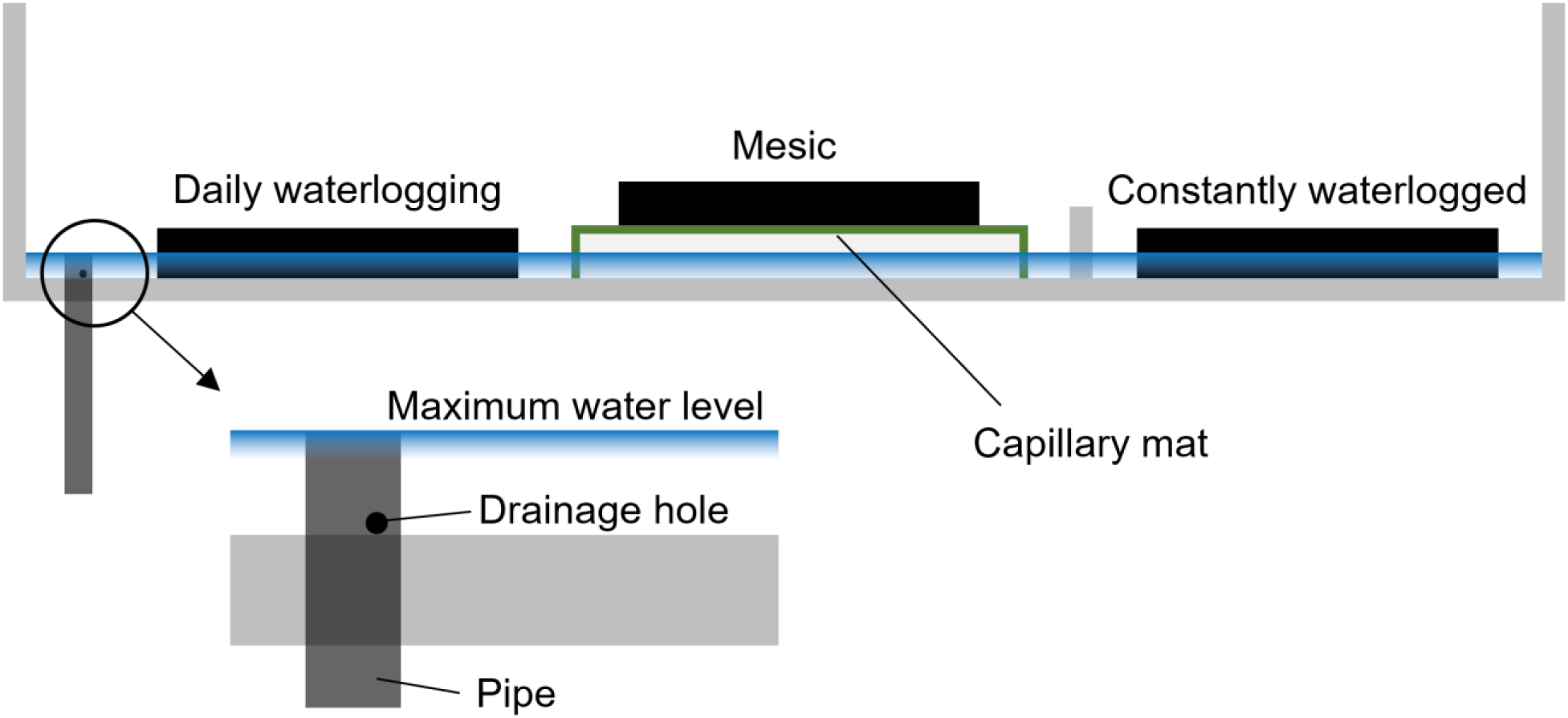
Experimental setup, with trays in black. Water was added daily, with the maximum water level controlled by the overflow into the pipe (open topped), the water level then dropped down to 0 over approximately 2 hours.

Before experimental treatments started, the samples were subjected to four weeks of cold, wet stratification at 4 °C. No seedlings germinated during this period. The samples were cultivated for 12 weeks, and within the first month 95% of the seedlings had germinated, and germination virtually ceased during the last month. The trays were checked twice a week, and large seedlings that could be identified correctly were removed. We followed the taxonomy of Krok & Almquist (2012) for identification. Seedlings were identified to the lowest possible taxonomic level (Supp. Table 1). Some specimens were identified to the level of genus because of a lack of identifiable organs (e.g., flowers in *Callitriche* spp). We found one seedling (*Juncus effusus*) in the control sections of a continuous waterlogging tray, which was likely a contamination from one of the neighboring samples rather than the growing media, as the species were present in both of those sites.

### Data analyses

To evaluate the effects of hydropeaking on riverine SSBs, we characterised SSBs by their density (i.e. total number of seedlings per sample), richness (i.e. number of species identified in a sample), their functional group (proportion of species typical for the graminoid belt, see Figure 1) and Ellenberg values (for Moisture, Light and Soil disturbance). For the latter, we used species ecological indicator values developed for Sweden (Tyler et al., 2021). The impact of hydropeaking on these characteristics was tested using generalized linear mixed-effect models (GLMMs) for the density, richness and proportion of large graminoids, taking treatment into account, and with site as a random factor. For the CWMs, we used a GLM, with the data summarized to site level. All analyses were conducted in R using RStudio (R version 4.4.2; RStudio Team, Boston, MA), using base R or package glmmTMB (Brooks et al., 2017). GLMMs were conducted with a nbinom2 (SSB richness) or a tweedie (SSB density and proportion large graminoids) distribution, and GLMs (CWMs) with a Gaussian distribution. Overdispersion and outliers in the residuals were checked with the help of the DHARMa package (Hartig, 2016), and were non-significant.

## Results

Across all treatments, 717 seedlings emerged during the germination experiment, of which 716 could be identified to one of 52 taxa (Supp. Table 1). The most common species were *Betula pendula/pubescens* (13 sites) and *Juncus bulbosus* (9 sites), and the most numerous was *Hypericum maculatum* (200 seedlings). On average, 8.8 seedlings of 3.0 species were detected per replicate, and 2 to 18 species per site. A mesic treated sample from Röjdan had the highest number of taxa (13) and shared the highest number of seedlings (47) with an Eldforsen sample under continuous waterlogging. No seedlings germinated from Långbjörn samples undergoing the daily waterlogging treatment.

### Seedling numbers and species richness

Samples from sites with stronger flow regulation tended to produce fewer seedlings than those of sites with milder regulation regimes (Figure 5). With a 1 cm increase in hourly change in water level, modelled seedling density decreased by 29% in the mesic treatment, 14% under daily waterlogging, and 9% under continuous waterlogging (Supp. Table 2). Seedling density did not significantly differ between treatments overall. We found no significant correlation between regulation intensity and seed bank species richness (Figure 6, for full model output see Supp. Table 3).

**Figure 5.**
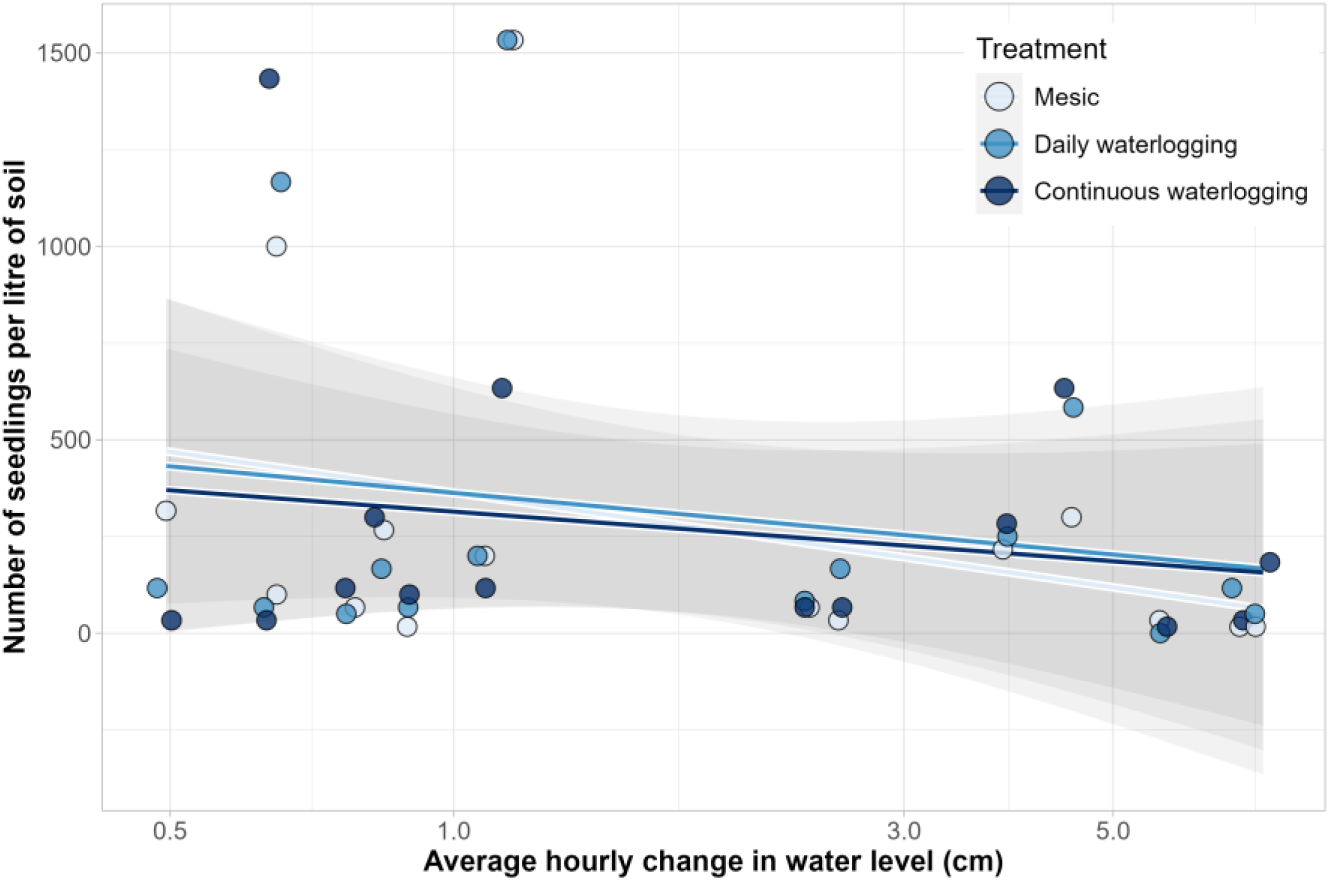
Seedling density relative to hydropeaking intensity (notice non-linear x-axis). Shading represents 95% confidence interval. Points slightly displaced for visibility.

**Figure 6.**
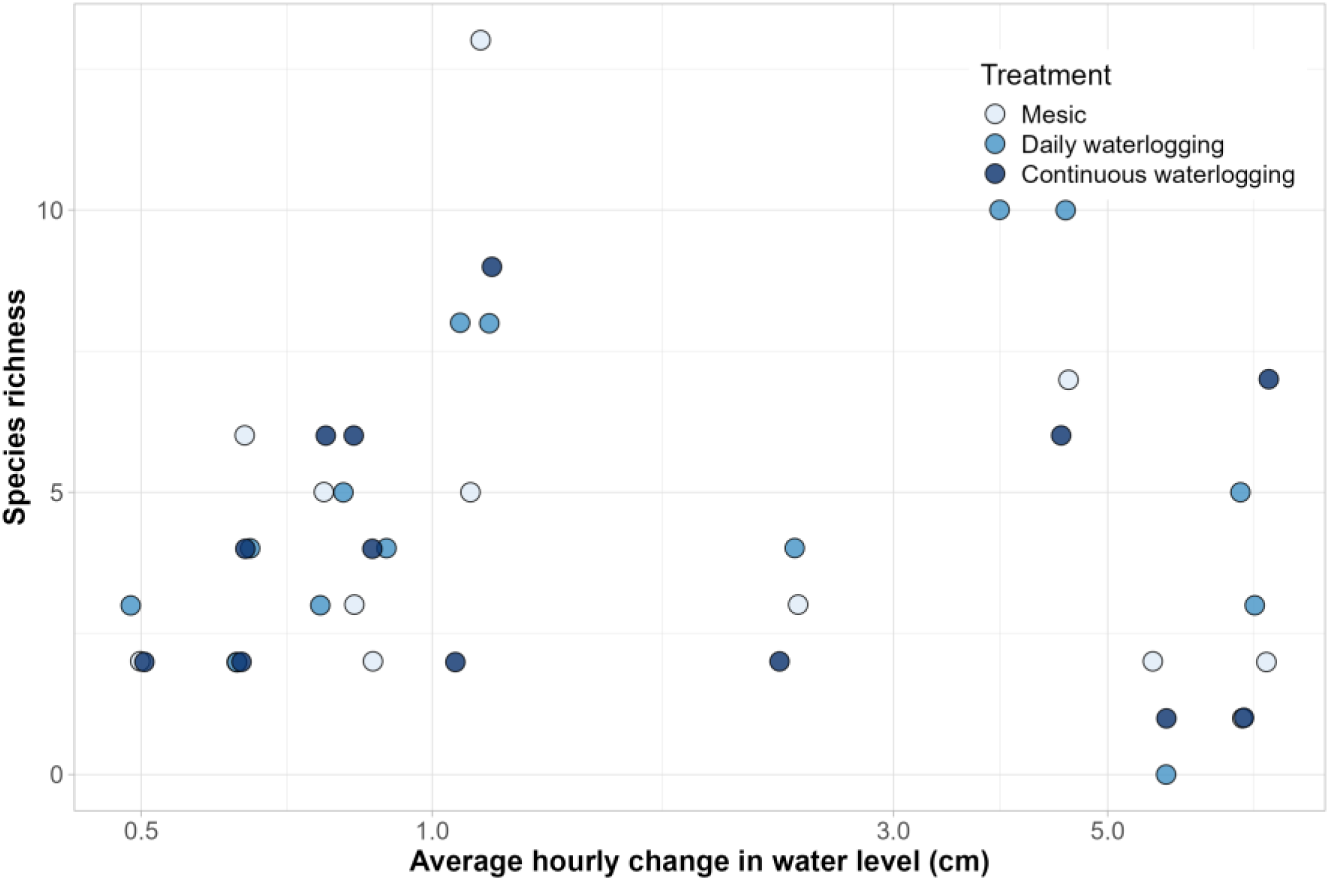
Species richness in relation to hydropeaking intensity (notice non-linear x-axis). Points slightly displaced for visibility.

### Functional responses

Large graminoids (Supp. Table 1), with typical riparian species such as *Carex acuta* and *Juncus effusus*, made up a larger proportion of the seedbank in samples from sites with relatively mild flow regulation (Supp. Table 4, Figure 7). The overall proportion decreased with flow regulation intensity, which translates to a statistically near-significant correlation and an overall 14.4% decrease per cm increase in hourly change. Community-weighted mean (CWM) values for Ellenberg light, moisture and soil disturbance showed a positive but non-significant correlation with increasing hydropeaking intensity (Supp. Table 5, Figure 8).

**Figure 7.**
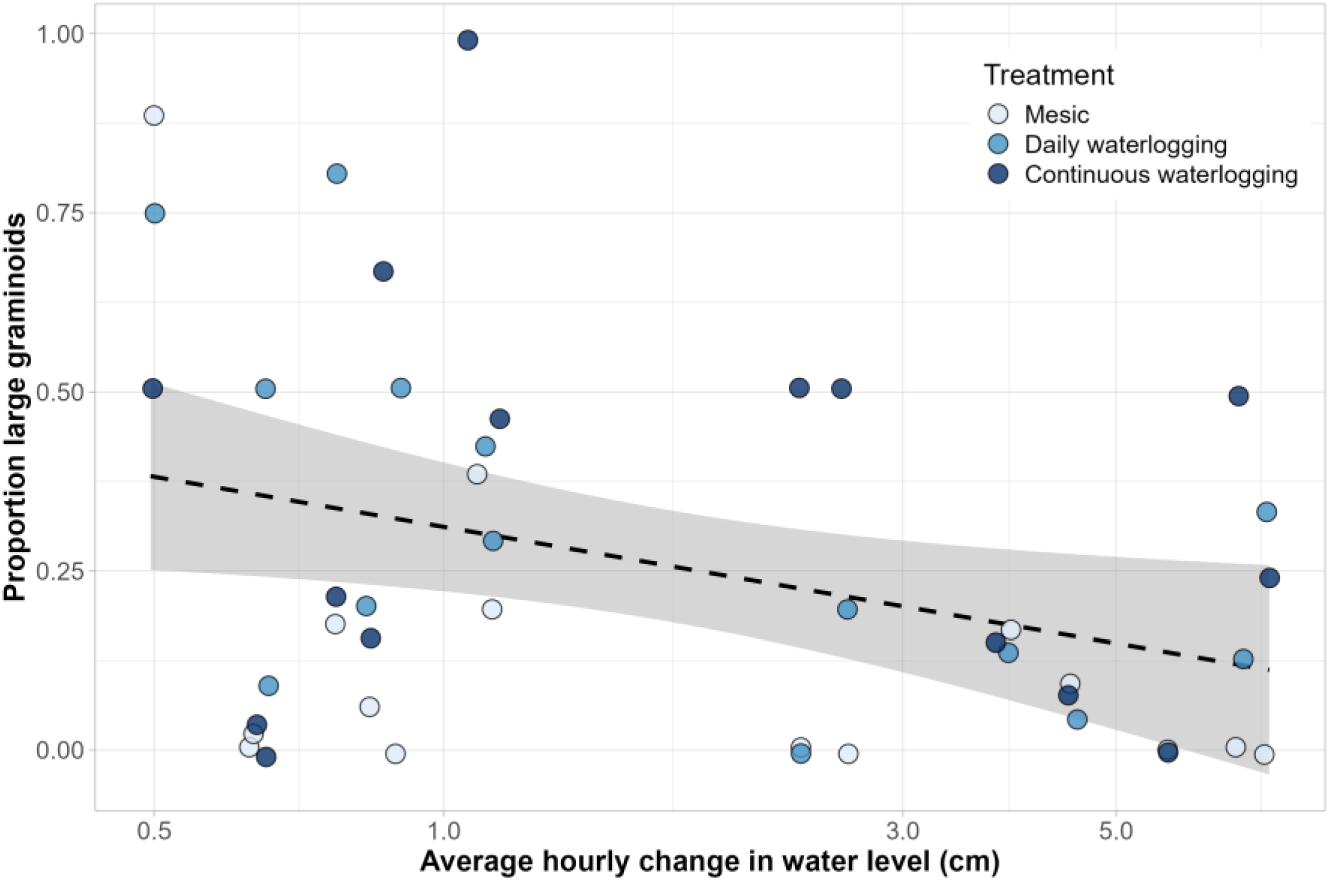
The proportion of large graminoids among seedlings against hydropeaking intensity (notice non-linear x-axis). Shading represents 95% confidence interval. Points slightly displaced for visibility.

**Figure 8.**
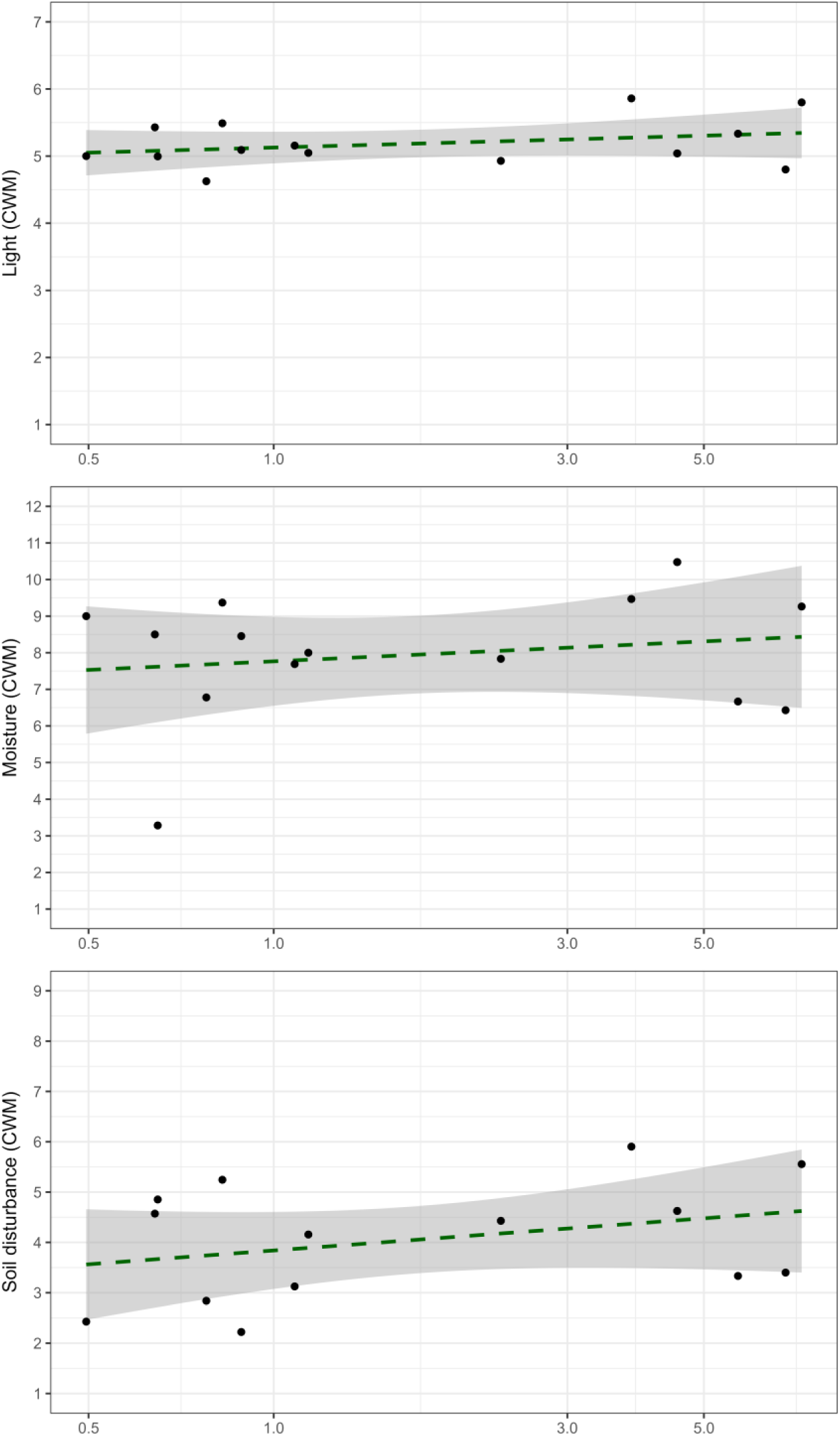
Community-weighted means of Ellenberg values calculated for each site (N = 14) against hydropeaking intensity (notice non-linear scale).

## Discussion

Through a seedling emergence experiment, we have tested whether and how riparian seed banks are affected by flow regulation. Riparian seed banks build through supply from local and regional (upstream) sources, and decrease through germination and die-off. Regulation results in altered hydrochory patterns, changed vegetation and thus local seed production, and different opportunities for seeds to germinate or remain in the seed bank (Andersson et al., 2000; Bejarano et al., 2020; Sarneel et al., 2016). In our germination experiment, we subjected seed bank samples from regulated rivers to three different treatments to account for different tolerances of seeds and thus get the most complete overview of the seed bank. We found that seed banks were relatively species-poor compared to riparian vegetation in the area (Hoppenreijs et al., submitted; Jansson et al., 2000). Seedling density was variable and weakly negatively correlated with regulation intensity. Especially the proportion of large graminoids in the seed bank decreased, while species tolerant of higher moisture and, especially, soil disturbance made up a somewhat larger proportion of seed banks of more heavily regulated sites.

### Regulation intensity and seed bank impoverishment

Seedling numbers were correlated negatively with flow regulation intensity, measured as average hourly change in water level (Figure 5). We used samples from relatively sheltered sites, where water is expected to deposit seeds. This may partly explain the weakness of the negative trend in seedling numbers in relation to hydropeaking intensity. Such a pattern might be stronger at more exposed sections where seed bank build-up would be hampered by both natural and hydropeaking caused erosion (Nordström et al., submitted). We know of few other studies of seed banks along regulated streams, but a South Australian example showed similar results with respect to the effect of regulation intensity on seedling numbers (Greet et al., 2013). The authors found that seedling numbers were higher along regulated rivers than along free-flowing rivers, mostly due to the large number of non-native seeds. We found no non-native seeds in our samples, similar to studies on riparian vegetation in the same area (Dynesius & Nilsson, 1994; Hoppenreijs et al., submitted), but made no comparison to unregulated rivers.

Although species might be affected differently by flow regulation, we did not see a decrease in species richness of SSBs with increasing regulation intensity (Figure 6). We did find high variation in species richness per site (minimum 2, median 8, maximum 18), for which there are multiple possible explanations. In the vegetation, naturally occurring species may be replaced by species that are better adapted to regulated environments (Hoppenreijs et al., submitted), or sites may experience an immediate decrease in species richness with regulation, irrespective of the degree of regulation (Jansson et al., 2000). Other factors, such as time since the regulation regime started (Su et al., 2025), distance from regulation source (Wollny et al., 2021), number of dams upstream, and contributions from tributaries may also play a role (Bång et al., 2007). The same factors may affect seed banks. Additionally, both limited dispersal by dams and increased erosion by changed flow and ice regimes may affect seed bank species richness negatively (Jansson et al., 2005; Nordström et al., submitted). Due to scale, these factors could not be taken into account in the current study, but the high variation in both seedling numbers and species richness in our dataset indicates the need for locality-specific information about the ecosystem, both to understand the risks of flow regulation and the potential for restoration.

### Species tolerances under different regulation intensities

We found that large graminoids are impacted negatively by increasing flow regulation (Figure 7, near-significant correlation). A belt of graminoids is common along free-flowing streams (e.g., Ström et al., 2012), and a decrease in the proportion of graminoids in the seed bank could thus create a shift in vegetation composition and is then likely to cause changed ecosystem functioning. Next to habitat provision to macroinvertebrates, large graminoids are for example important for erosion prevention (Micheli & Kirchner, 2002).

In the trays undergoing the mesic treatment, i.e. daily watering and drying up after some hours, we found decreasing germination with increasing regulation intensity. This suggests that there are fewer species that respond well to drier conditions left in SSBs of heavily regulated streams, which is confirmed by the community-weighted Ellenberg values (Figure 8). If riparian seed banks are largely a product of local seed production, since hydrochory is limited in regulated systems (Andersson et al., 2000), this can be explained by decreased abundance and reproductive success of riparian plants that are intolerant of frequent and prolonged flooding (Bejarano et al., 2018, 2020).

Together, the effect on graminoids and mesic species could decrease the zonation of the riparian zone, and possibly leave areas open to invasion by species detrimental to the function of the riparian zones (such as ruderals or non-natives as seen by Greet et al., 2013). The Ellenberg values are possibly pointing towards such a shift, with slightly higher values for light and soil disturbance (Figure 8) at sites with higher hydropeaking intensities. As seed banks are slower to respond to change than standing vegetation (Goodson et al., 2001), there is also a possibility that this effect will grow more pronounced over time, and restoration may be more successful the sooner it is attempted.

### Adapted flow management and riparian restoration

While flow regulation remains an important source of renewable energy, research shows that small adaptations in flow management can lead to relatively large ecological benefits (Widén et al., 2022). If such adaptations are made, seed banks may be an early source of recovery for riparian vegetation, especially when communities are dispersal-limited (Hasselquist et al., 2015; Sarneel et al., 2024). However, with the rather poor seed banks like the ones we found in our system, seed addition or even planting may be a measure to kickstart the recovery of the system. Seeding may be especially helpful in frequently-disturbed habitats due to their invasion-proneness, or when the seed rain is expected to be of low quality (Florentine et al., 2023; Wells et al., 2024). As diversity in growth forms is indicative of riparian zone condition (O’Donnell et al., 2016), seeding of species or groups known to be negatively affected by earlier regulation regimes could help steer the recovery process.

## Data Availability Statement

The data is available from the corresponding author upon reasonable request.

## Acknowledgements

We thank Jennifer Lehikoinen for help with fieldwork, Geni Zanol for help with the growing room, and Oskar Gedda and Susanne Larsson for their help with the germination experiment and for providing input to an earlier version of the manuscript. We thank Lutz Eckstein for his help during the conceptualisation of the study.

## Supplementary materials

**Table 1.**
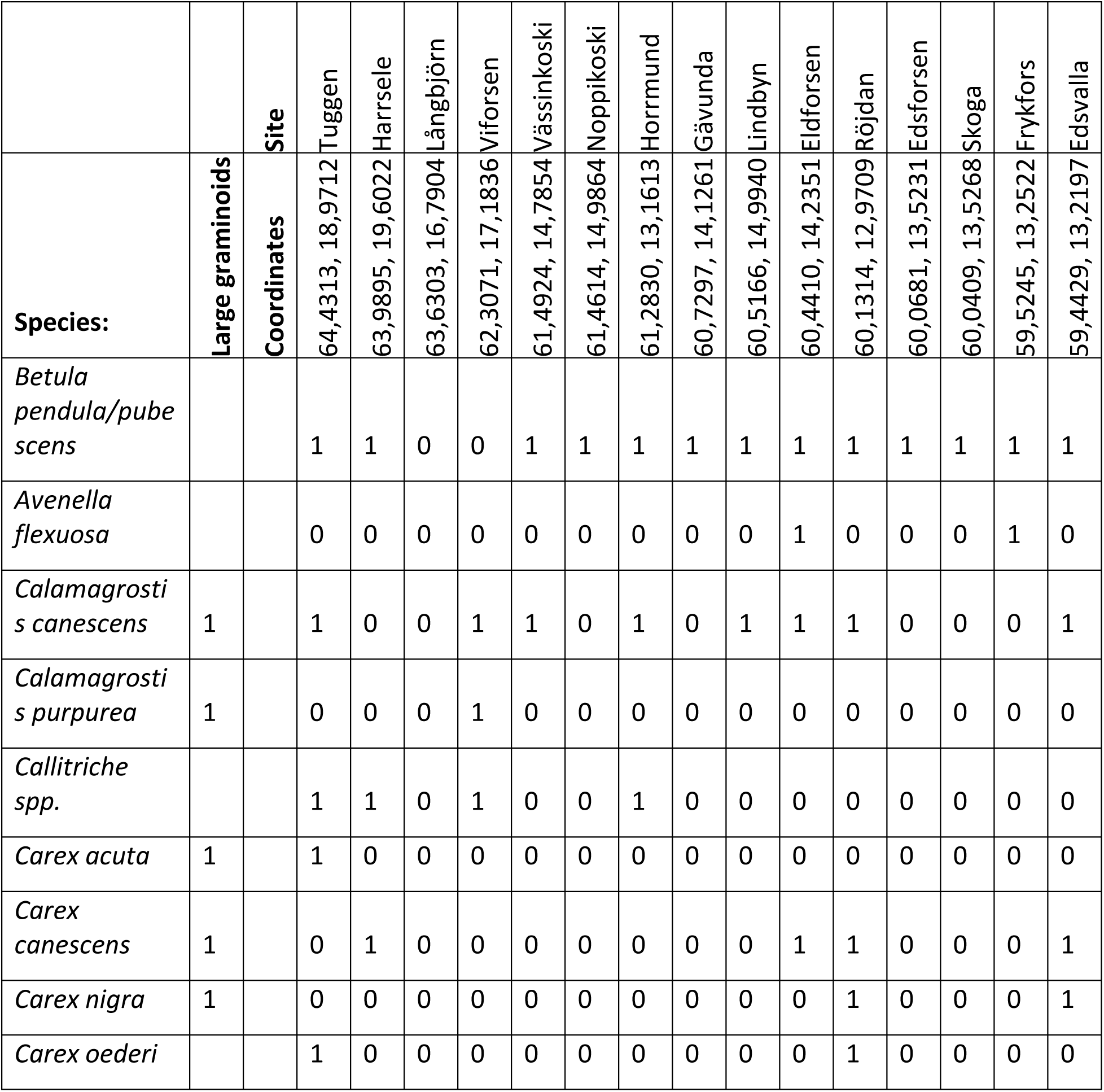
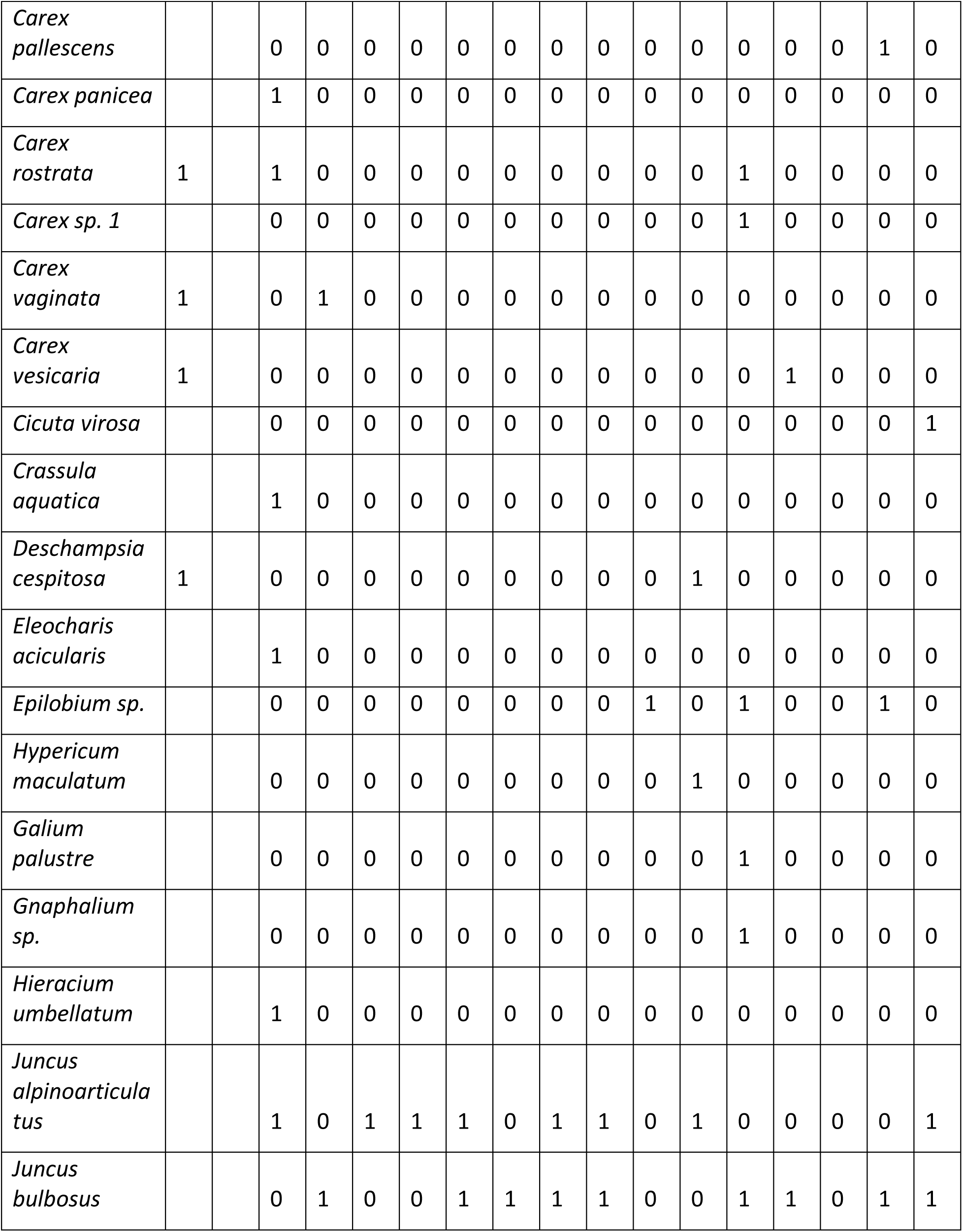
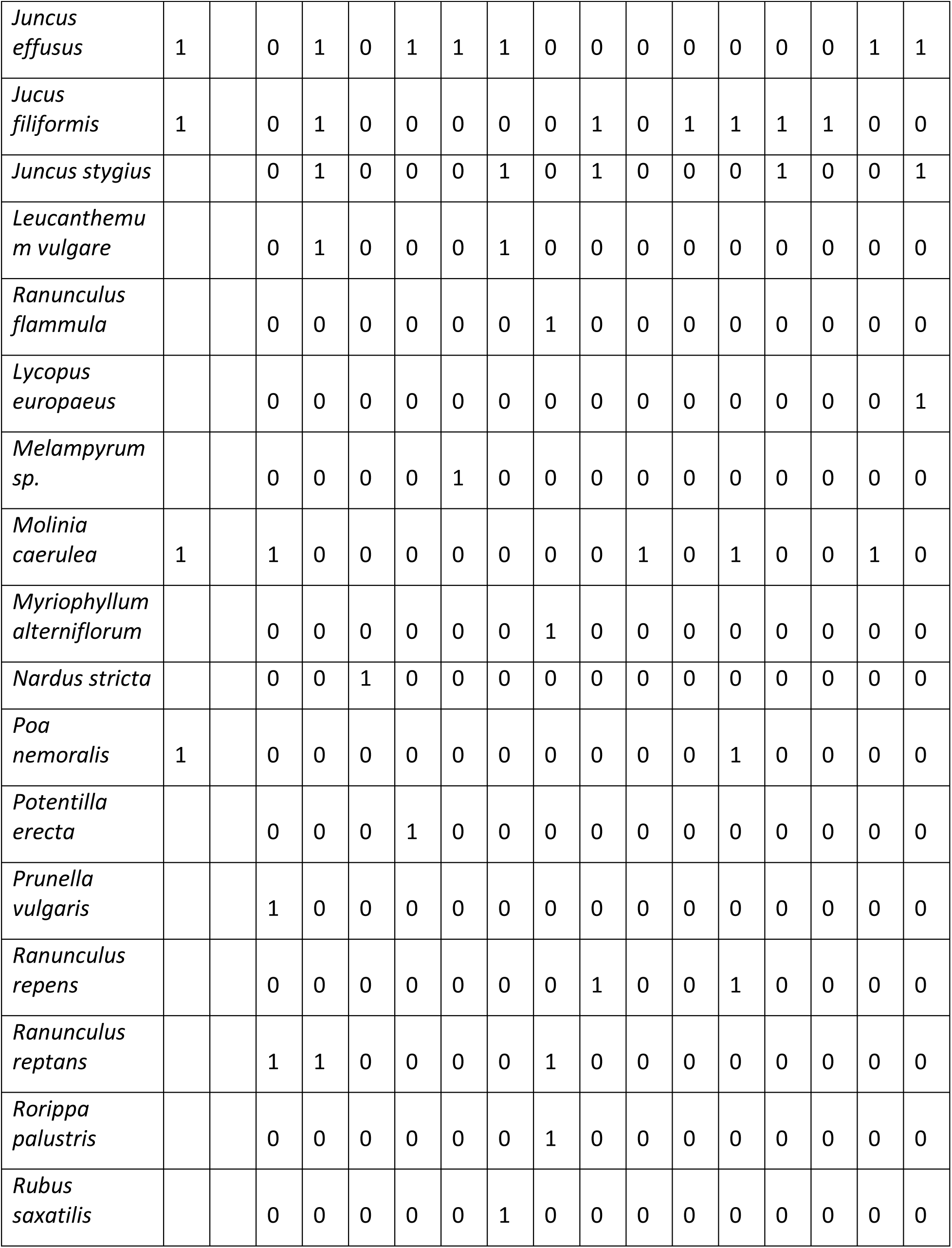
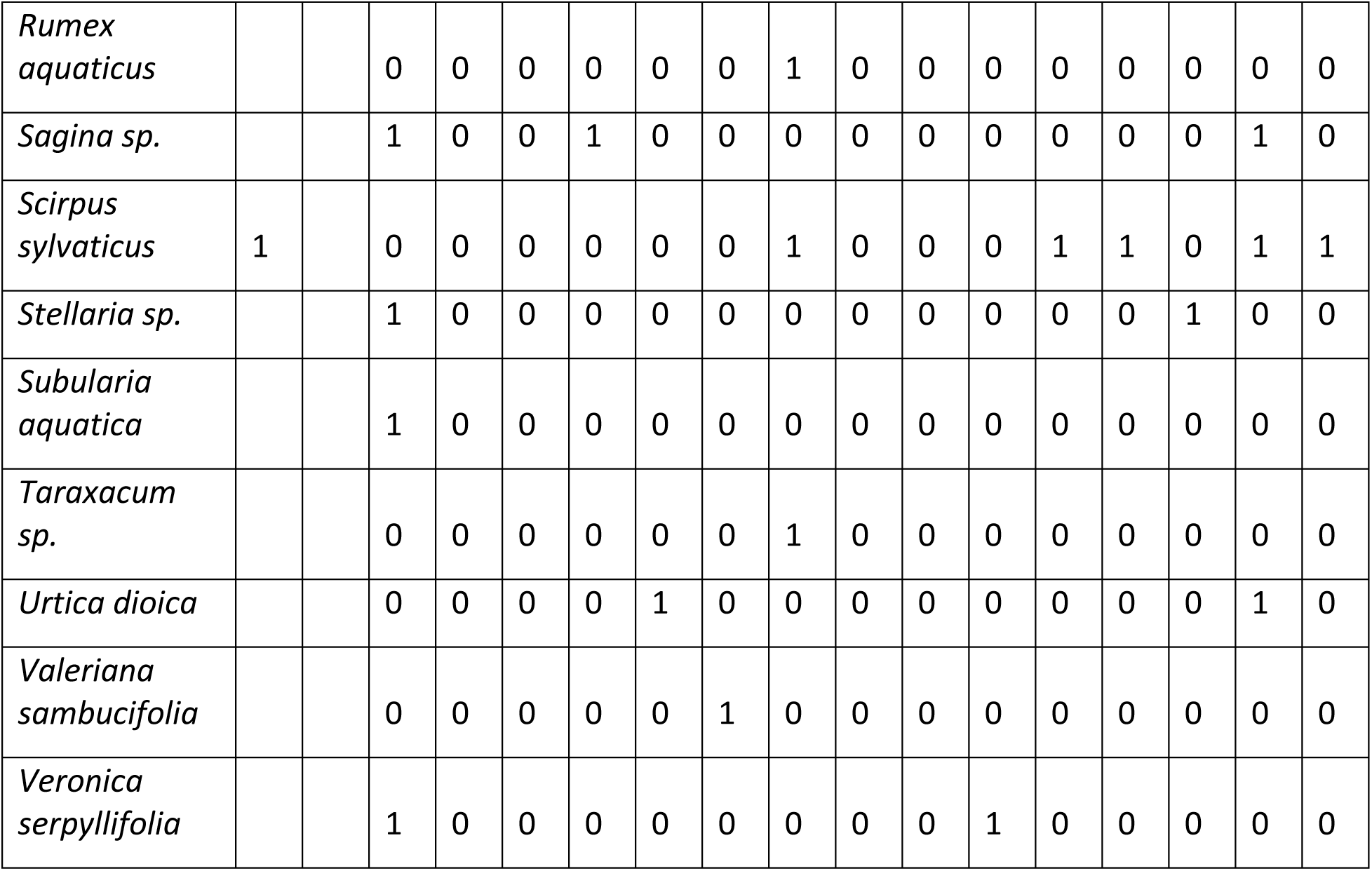
seed bank species list. Species list, with large graminoids marked, and at which site(s) the species occurred (arranged from north to south).

**Table 2.** GLMM output seedling counts. Mesic as reference value, p values <0.05 in bold.

| Term | Estimate | Std. Error | z value | Pr(>\|z\|) |
| --- | --- | --- | --- | --- |
| (Intercept) | 2.1943 | 0.4421 | 4.9638 | **<0.0001** |
| Hydropeaking intensity (cm/h) | -0.3352 | 0.1375 | -2.4381 | **0.0148** |
| Treatment(Daily waterlogging) | -0.2261 | 0.2338 | -0.9671 | 0.3335 |
| Treatment(Continuous waterlogging) | -0.4471 | 0.2380 | -1.8787 | 0.0603 |
| Hydropeaking intensity X Treatment(Daily waterlogging) | 0.1832 | 0.0920 | 1.9916 | **0.0464** |
| Hydropeaking intensity X Treatment(Continuous waterlogging) | 0.2405 | 0.0911 | 2.6393 | **0.0083** |

**Table 3.**
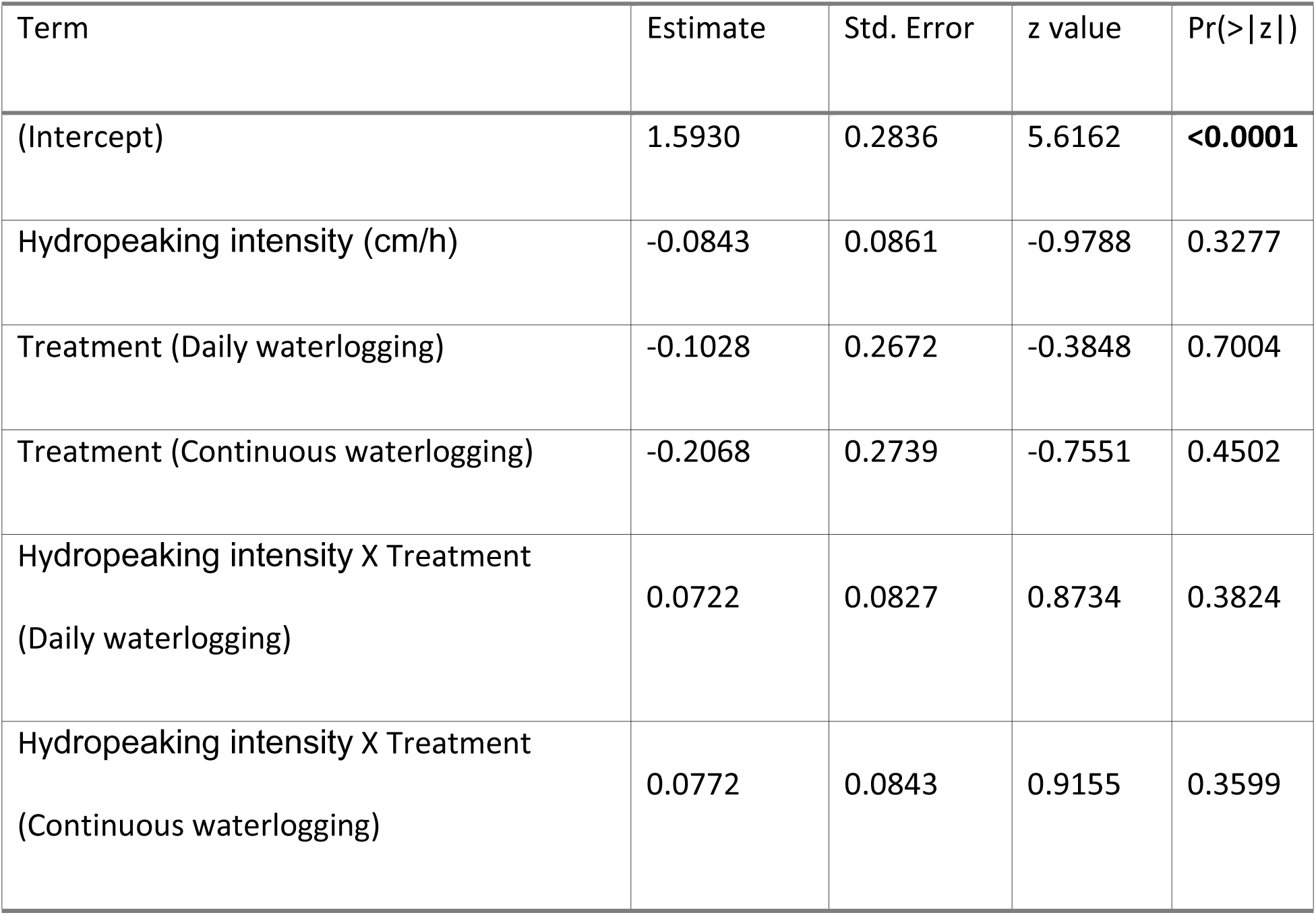
GLMM output of species richness. Mesic as reference value, p values <0.05 in bold.

**Table 4.**
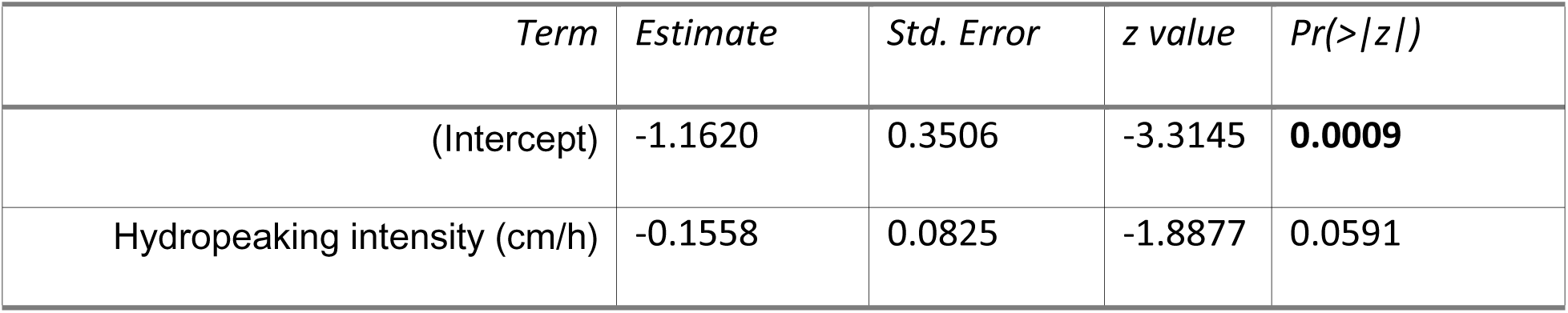
GLMM output of proportion large graminoids.

**Table 5.** GLM output of CWMs for light, moisture and soil disturbance.

| Trait | Term | Estimate | Std. Error | t value | Pr(>\|t\|) |
| --- | --- | --- | --- | --- | --- |
| Light | (Intercept) | 0.6879 | 0.0181 | 38.1004 | **<0.0001** |
|  | log(Hydropeaking intensity) | 0.0183 | 0.0167 | 1.1000 | 0.2929 |
| Moisture | (Intercept) | 0.6150 | 0.0507 | 12.1293 | **<0.0001** |
|  | log(Hydropeaking intensity) | 0.0308 | 0.0468 | 0.6571 | 0.5235 |
| Soil<br>disturbance | (Intercept) | 0.3550 | 0.0439 | 8.0875 | **<0.0001** |
|  | log(Hydropeaking intensity) | 0.0496 | 0.0406 | 1.2226 | 0.2449 |

## References

Andersson, E., Nilsson, C., & Johansson, M. E. (2000). Effects of river fragmentation on plant dispersal and riparian flora. River Research and Applications, 16(1), 83–89. 10.1002/(sici)1099-1646(200001/02)16:1%3C83::aid-rrr567%3E3.0.co;2-t

Arthington, A. H., Naiman, R. J., McClain, M. E., & Nilsson, C. (2010). Preserving the biodiversity and ecological services of rivers: New challenges and research opportunities. Freshwater Biology, 55, 1–16. 10.1111/j.1365-2427.2009.02340.x

Baird, I. G., Ziegler, A. D., Fearnside, P. M., Pineda, A., Sasges, G., Strube, J., Thomas, K. A., Schmutz, S., Greimel, F., & Hayes, D. S. (2024). Ruin-of-the-rivers? A global review of run-of-the-river dams. Environmental Management. 10.1007/s00267-024-02062-5

Baker, D. B., Richards, R. P., Loftus, T. T., & Kramer, J. W. (2004). A New Flashiness Index: Characteristics and Applications to Midwestern Rivers and Streams. Journal of the American Water Resources Association, 40(2), 503–522. 10.1111/j.1752-1688.2004.tb01046.x

Bejarano, M. D., Jansson, R., & Nilsson, C. (2018). The effects of hydropeaking on riverine plants: A review. Biological Reviews, 93(1), 658–673. 10.1111/brv.12362

Bejarano, M. D., Sordo-Ward, Á., Alonso, C., Jansson, R., & Nilsson, C. (2020). Hydropeaking affects germination and establishment of riverbank vegetation. Ecological Applications, 30(4), 1–16. 10.1002/eap.2076

Brooks, M., Bolker, B., Kristensen, K., Maechler, M., Magnusson, A., Skaug, H., Nielsen, A., Berg, C., & Van Bentham, K. (2017). glmmTMB: Generalized Linear Mixed Models using Template Model Builder (p. 1.1.14) [Data set]. 10.32614/CRAN.package.glmmTMB

Bång, Å., Nilsson, C., & Holm, S. (2007). The potential role of tributaries as seed sources to an impoundment in Northern Sweden: A field experiment with seed mimics. River Research and Applications, 23(10), 1049–1057. 10.1002/rra.1014

Corenblit, D., Tabacchi, E., Steiger, J., & Gurnell, A. M. (2007). Reciprocal interactions and adjustments between fluvial landforms and vegetation dynamics in river corridors: A review of complementary approaches. Earth-Science Reviews, 84(1–2), 56–86. 10.1016/j.earscirev.2007.05.004

Dynesius, M., & Nilsson, C. (1994). Fragmentation and flow regulation of river systems in the northern third of the world. Science, 266(5186), 753–762. 10.1126/science.266.5186.753

Dudgeon, D. (2019). Multiple threats imperil freshwater biodiversity in the Anthropocene. Current Biology, 29(19), R960–R967. 10.1016/j.cub.2019.08.002

Florentine, S., Milberg, P., & Westbrooke, M. (2023). Potential contributions of the soil seed bank and seed rain for accelerating the restoration of riparian catchments in Australia. In Global Ecology and Conservation, (Vol. 47). ELSEVIER. 10.1016/j.gecco.2023.e02645

Greet, J., Cousens, R. D., & Webb, J. A. (2013). Flow regulation is associated with riverine soil seed bank composition within an agricultural landscape: Potential implications for restoration. Journal of Vegetation Science, 24, 157–167. 10.1111/j.1654-1103.2012.01445.x

Hall, S. J., & Zedler, J. B. (2010). Constraints on Sedge Meadow Self-Restoration in Urban Wetlands. Restoration Ecology, 18(5), 671–680. 10.1111/j.1526-100X.2008.00498.x

Harris, H. A. L., Murray, T. J., Tonkin, J. D., & McIntosh, A. R. (2026). Asynchronous river floodplain environment dampens ecological variability across scales. Oikos, e11967. 10.1002/oik.11967

Hartig, F. (2016). DHARMa: Residual Diagnostics for Hierarchical (Multi-Level / Mixed) Regression Models (p. 0.4.7) [Data set]. 10.32614/CRAN.package.DHARMa

Hasselquist, E. M., Nilsson, C., Hjältén, J., Jørgensen, D., Lind, L., & Polvi, L. E. (2015). Time for recovery of riparian plants in restored northern Swedish streams: A chronosequence study. Ecological Applications, 25(5). 10.1890/14-1102.1

He, F., Zarfl, C., Tockner, K., Olden, J. D., Campos, Z., Muniz, F., Svenning, J.-C., & Jähnig, S. C. (2024). Hydropower impacts on riverine biodiversity. Nature Reviews Earth & Environment. 10.1038/s43017-024-00596-0

Hoppenreijs, J. H. T., Eckstein, R. L., & Lind, L. (2022). Pressures on boreal riparian vegetation: A literature review. Frontiers in Ecology and Evolution, 9, Article 806130. 10.3389/fevo.2021.806130

Hoppenreijs, J. H. T., Eckstein, R. L., & Lind, L. (submitted). Flow regulation effects on riparian width and vegetation composition: not fewer, but different species. River Research and Applications.

Jansson, R., Nilsson, C., Dynesius, M., & Andersson, E. (2000). Effects of river regulation on river-margin vegetation: A comparison of eight boreal rivers. Ecological Applications, 10(1), 203–224. 10.1890/1051-0761(2000)010%5B0203:EORROR%5D2.0.CO;2

Jansson, R., Ström, L., & Nilsson, C. (2019). Smaller future floods imply less habitat for riparian plants along a boreal river. Ecological Applications, 29(8), Article e01977. 10.1002/eap.1977

Jansson, R., Zinko, U., Merritt, D. M., & Nilsson, C. (2005). Hydrochory increases riparian plant species richness: A comparison between a free-flowing and a regulated river. Journal of Ecology, 93(6), 1094–1103. 10.1111/j.1365-2745.2005.01057.x

Krok, Th. O. B. N., & Almquist, S. (2013). Svensk flora: Fanerogamer och kärlkryptogamer (L. Jonsell & B. Jonsell, Eds). Liber.

Lozanovska, I., Rivaes, R., Vieira, C., Ferreira, M. T., & Aguiar, F. C. (2020). Streamflow regulation effects in the Mediterranean rivers: How far and to what extent are aquatic and riparian communities affected? Science of The Total Environment, 749, 141616. 10.1016/j.scitotenv.2020.141616

Lytle, D. A., Merritt, D. M., Tonkin, J. D., Olden, J. D., & Reynolds, L. V. (2017). Linking river flow regimes to riparian plant guilds: A community-wide modeling approach. Ecological Applications, 27(4). 10.1002/eap.1528

Micheli, E. R., & Kirchner, J. W. (2002). Effects of wet meadow riparian vegetation on streambank erosion. 2. Measurements of vegetated bank strength and consequences for failure mechanics. Earth Surface Processes and Landforms, 27(7), 687–697. 10.1002/esp.340

Nakano, S., & Murakami, M. (2001). Reciprocal subsidies: Dynamic interdependence between terrestrial and aquatic food webs. Proceedings of the National Academy of Sciences of the United States of America, 98(1), 166–170. 10.1073/pnas.98.1.166

Nordström, E., Watz, J., Greenberg, L., Malm Renöfält, B., Jansson, R., Eckstein, R. L., & Bergman, E. (submitted). Stream ice dynamics and riparian disturbance under hydropeaking flows. Journal of Hydrology.

Nyqvist, D., Calles, O., Carlson, P., Holmgren, K., Malm-Renöfält, B., Widén, Å., Bergengren, J., & Näslund, J. (2025). Balancing hydropower production and ecology − ecological impacts, mitigation measures, and programmatic monitoring. Knowledge & Management of Aquatic Ecosystems, 426(24).

O’Donnell, J., Fryirs, K. A., & Leishman, M. R. (2016). Seed banks as a source of vegetation regeneration to support the recovery of degraded rivers: A comparison of river reaches of varying condition. Science of the Total Environment, 542, 591–602. 10.1016/j.scitotenv.2015.10.118

O’Donnell, J., Fryirs, K., & Leishman, M. R. (2015). Can the regeneration of vegetation from riparian seed banks support biogeomorphic succession and the geomorphic recovery of degraded river channels? River Research and Applications, 31(7), 834–846. 10.1002/rra.2778

Sarneel, J. M., Brett, L., Van Den Bosch-Hennink, A., & Hasselquist, E. M. (2024). River restoration effects on dispersal and the development of riparian seed bank: Do poor seed banks limit restoration of boreal riparian zones? Restoration Ecology, e14328. 10.1111/rec.14328

Sarneel, J. M., Kardol, P., & Nilsson, C. (2016). The importance of priority effects for riparian plant community dynamics. Journal of Vegetation Science, 27(4), 658–667. 10.1111/jvs.12412

Ström, L., Jansson, R., & Nilsson, C. (2012). Projected changes in plant species richness and extent of riparian vegetation belts as a result of climate-driven hydrological change along the Vindel River in Sweden: Climate-change effects on riparian vegetation. Freshwater Biology, 57(1), 49–60. 10.1111/j.1365-2427.2011.02694.x

Su, X., Bejarano, M. D., Jansson, R., Pilotto, F., Sarneel, J. M., Lin, F., Wang, Y., Cai, F., Wu, S., & Zeng, B. (2025). Broad-Scale Meta-Analysis of Drivers Mediating Adverse Impacts of Flow Regulation on Riparian Vegetation. Global Change Biology, 31(2), e70042. 10.1111/gcb.70042

Ter Heerdt, G. N. J., Verweij, G. L., Bekker, R. M., & Bakker, J. P. (1996). An improved method for seed-bank analysis: Seedling emergence after removing the soil by sieving. Functional Ecology, 10(1), 144–151. 10.2307/2390273

Tyler, T., Herbertsson, L., Olofsson, J., & Olsson, P. A. (2021). Ecological indicator and traits values for Swedish vascular plants. Ecological Indicators, 120, Article 106923. 10.1016/j.ecolind.2020.106923

Ward, J. V. (1989). The Four-Dimensional Nature of Lotic Ecosystems. Journal of the North American Benthological Society, 8(1), 2–8.

Wells, A. J., Harrington, J., & Balster, N. J. (2024). Seeding Density Alters the Assembly of a Restored Plant Community after the Removal of a Dam in Southern Wisconsin, USA. Environments, 11(6), 115. 10.3390/environments11060115

Widén, Å., Malm Renöfält, B., Degerman, E., Wisaeus, D., & Jansson, R. (2022). Environmental flow scenarios for a regulated river system: Projecting catchment-wide ecosystem benefits and consequences for hydroelectric production. Water Resources Research, 58(1), Article e2021WR030297. 10.1029/2021WR030297

Wollny, J. T., Bergmann, W., Otte, A., & Harvolk-Schöning, S. (2021). River regulation intensity matters: Riverbank vegetation is characterized by more typical riverbank plant species with increasing distance from weirs. Ecological Engineering, 159, Article 106082. 10.1016/j.ecoleng.2020.106082

